# Photosynthesis and crop productivity is enhanced by glucose-functionalized fluorescent carbon dots

**DOI:** 10.1101/826628

**Authors:** Thomas A. Swift, Daniel Fagan, David Benito-Alifonso, Stephen A. Hill, Marian L. Yallop, Thomas A. A. Oliver, Tracy Lawson, M. Carmen Galan, Heather M. Whitney

## Abstract

From global food security to textile production and biofuels, the demands currently made on plant photosynthetic productivity will continue to increase. Enhancing photosynthesis using designer, green and sustainable materials offers an attractive alternative to current genetic-based strategies and promising work with nanomaterials has recently started to emerge. Here we describe *in planta* use of carbon-based nanoparticles produced by low-cost renewable routes that are bioavailable to mature plants. Uptake of these functionalised nanoparticles from the soil improves photosynthesis and also increases crop production. We show for the first time that glucose-functionalization enhances nanoparticle uptake, photoprotection and pigment production, unlocking enhanced yields. This is demonstrated in *Triticum aestivum* ‘Apogee’ (dwarf bread wheat) and results in an 18% increase in grain yield. This establishes the viability of a functional nanomaterial to augment photosynthesis as a route to increased crop productivity.

## INTRODUCTION

Plants greatly underperform compared to the theoretical maximum photosynthetic efficiency.^1,2^ Even under moderate solar light intensities, plants often absorb light in excess of that which can be safely harnessed due to downstream rate-limiting electron transport. Consequentially, plants take advantage of an inbuilt suite of photoprotective mechanisms, collectively termed non-photochemical quenching (NPQ), which dissipates excess energy and prevents deleterious photochemical reactions.^3^ Whilst essential to plant survival, it is widely accepted that NPQ impacts the photosynthetic performance of many plant species, with recent studies demonstrating that transgenic modification that targeted reducing specific components of NPQ, or downstream molecular and enzymatic processes, resulted a 15% increase in crop biomass.^4,5^ While recent work has highlighted that nanoparticles (NPs) offer a promising method for enhancing photosynthesis, these studies have yet to achieve the significant desired increases in crop yield.^6,7^

The work presented here accomplishes a dramatic increase in crop productivity by augmenting plants with specially engineered NPs. Here, the use of carbon dots (CDs) is explored to overcome the limitations of previously used nanomaterials such as toxicity, poor bioavailability or complex and inefficient syntheses.^6^ CDs are a carbon-based, fluorescent, small (less than 10 nm) class of nanoparticle.^8–11^ Our team and others have previously demonstrated that glycan-surface functionalization of NPs mitigates acute toxicity and enhances internalization in mammalian cells.^11–15^

Herein we report the first example of glycan-functionalized CDs used to significantly increase grain yields of elite bread wheat by 18%. Our mechanistic investigations reveal that CD-photosynthetic interactions affects many facets of photosynthesis, and that glycan-functionalization of CDs is essential to realize improved yields by altering NPQ and pigment production whilst also reducing reactive oxygen species generation.

## RESULTS

### Synthesis of functionalized fluorescent CDs

Unfunctionalized amine-coated CDs (core-CDs) were synthesized in one-pot after three-minutes microwave heating of glucosamine.HCl and 4,7,10-Trioxa-1,13-tridecanediamine following a modified reported procedure.^11,14^ Glucose-functionalization of the CDs was carried out in a two-step process by reaction of core-CDs with succinic anhydride to yield carboxylic acid bearing CDs, followed by N-(3-Dimethylaminopropyl)-N′-ethylcarbodiimide hydrochloride (EDC)-mediated conjugation of 1-aminoglucose to generate glucose-functionalized CDs (glucose-CDs) (Scheme S1, Figures S1-S6).

### CDs uptake in plants

The mechanisms by which the CDs interact with plants were then investigated in *Triticum aestivum* treated with core-CDs or glucose-CDs and compared to untreated controls. Each of these CDs were applied directly to the soil from three weeks post germination. CD uptake from the soil was observed by fluorescence microscopy for both CD treatments, (Figure 1 and S7) which have a peak fluorescence intensity at 455 nm,^11^ and are spectrally isolated from chlorophyll emission. The CD uptake was quantified, yielding 29 ± 2 µg and 32 ± 1 µg per gram of leaf tissue for core- and glucose-CDs, respectively. This measurement demonstrates that glucose-functionalization slightly enhances plant CD uptake (*p* ≤ 0.001, SI.)

**Figure 1.**
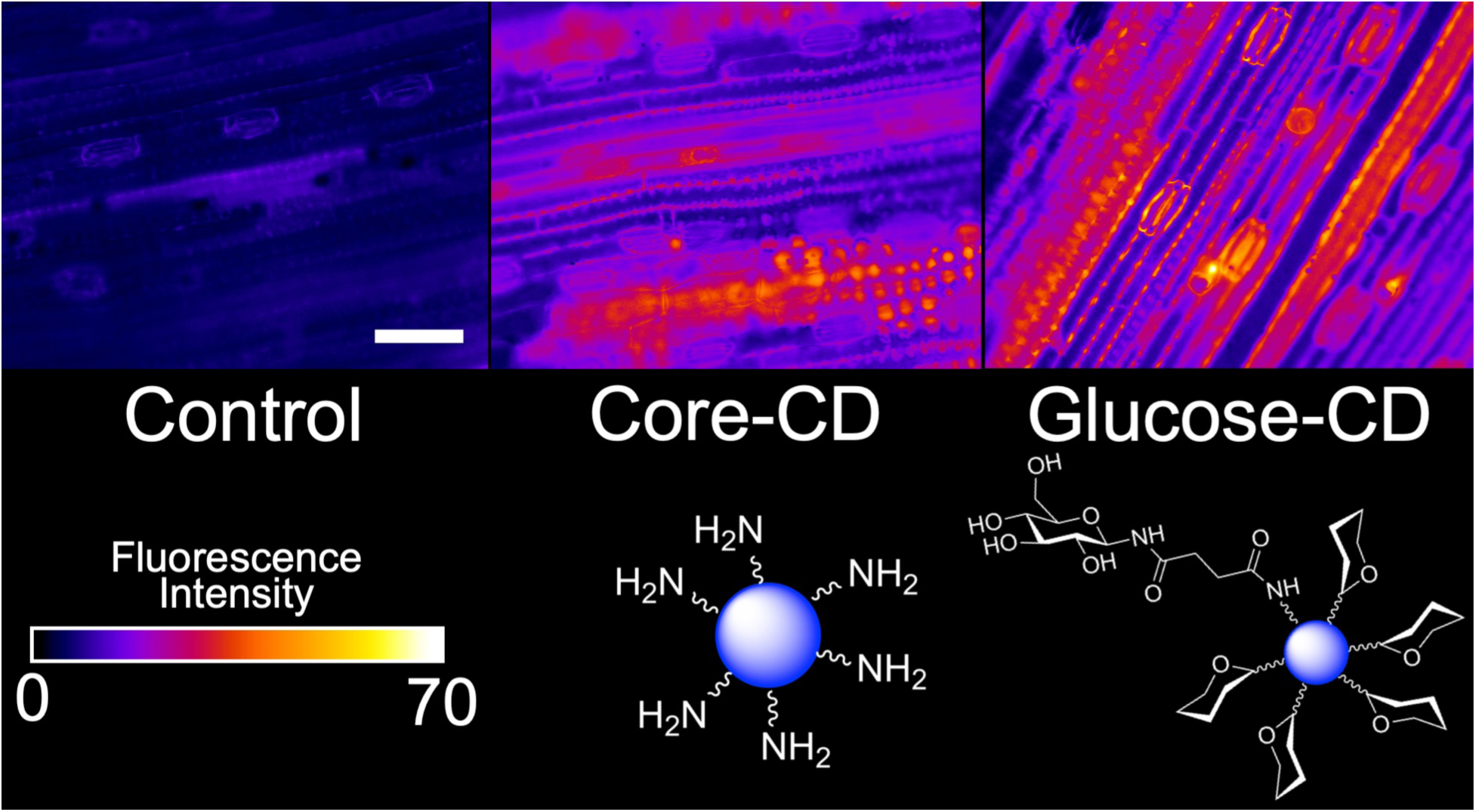
Fluorescence microscopy of CDs in *Triticum aestivum* mesophyll. Tissues were taken from the fully expanded flag leaves. An increase in fluorescence intensity was observed for the CD treatments (core- and glucose-CD) when compared to untreated controls. All images are the displayed on the same scale, with the white scale bar representing 100 μm. Fluorescence intensities are also displayed on the same scale. Schematics of the structure of the NP treatments are simplified and not to scale. Images were collected with 365 nm excitation (resonant with CD) and 455 nm emission.

### Effect of CDs on photosynthetic performance in plants

The effect of the CD-treatments on the photosynthetic performance of *Triticum aestivum* was investigated using standard protocols which measure chlorophyll fluorescence and infrared gas analysis (IRGA) as a function of photosynthetic photon flux densities (PPFD). At high actinic-light intensities, a small but significant enhancement of the operating efficiency of Photosystem II (*F*_*q*_’/*F*_*m*_’) was observed for both CD treatments (Figure 2, A) compared to the control. Furthermore, we observed a significant decrease in NPQ for plants treated with glucose-CDs compared to the other samples (Figure 2, B). Moreover, both CD treatments resulted in a significant increase of carbon assimilation (*A*) and the stomatal conductance (rate of CO_2_ passage through stomata, *g*_*s*_) compared to control (*p* < 0.05) for a wide range of PPFD (Figure 2, C and D). The *A* under saturating light conditions (*A*_*SAT*_) and the maximum *A* (*A*_MAX_) were extracted from exponential fits to data in Figure 2,C and S9, respectively. The values for *A*_*SAT*_ and *A*_*MAX*_ for each treatment are given in Table 1, with *A*_MAX_ invariant to CD treatment, but *A*_SAT_ enhanced by CD treatment.

**Table 1.** Calculated photosynthetic parameters.

| Parameter | Treatment |  |  |
| --- | --- | --- | --- |
|  | Control | Core-CD | Glucose-CD |
| $A_{SAT}$ ($\mu\text{mol m}^{-2} \text{s}^{-1}$) | $22.0 \pm 0.8$ | $24.0 \pm 0.9$ | $24.0 \pm 0.9$ |
| $A_{MAX}$ ($\mu\text{mol m}^{-2} \text{s}^{-1}$) | $31 \pm 2$ | $31 \pm 1$ | $30 \pm 1$ |
| $SI_{MAX}$ ($\mu\text{mol m}^{-2} \text{s}^{-2}$) | $222 \pm 6$ | $336 \pm 4$ | $244 \pm 7$ |
| Amplitude of NPQ<sub>1</sub> | $0.574 \pm 0.008$ | $0.50 \pm 0.01$ | $0.71 \pm 0.01$ |
| $\tau_1$ (s) | $85 \pm 3$ | $72 \pm 5$ | $94 \pm 3$ |
| Amplitude of NPQ<sub>2</sub> | $0.42 \pm 0.04$ | $0.502 \pm 0.004$ | $0.29 \pm 0.03$ |
| $\tau_2$ ($\times 10^3$ s) | $7 \pm 2$ | $10.3 \pm 0.4$ | $3.0 \pm 0.6$ |

**Figure 2.**
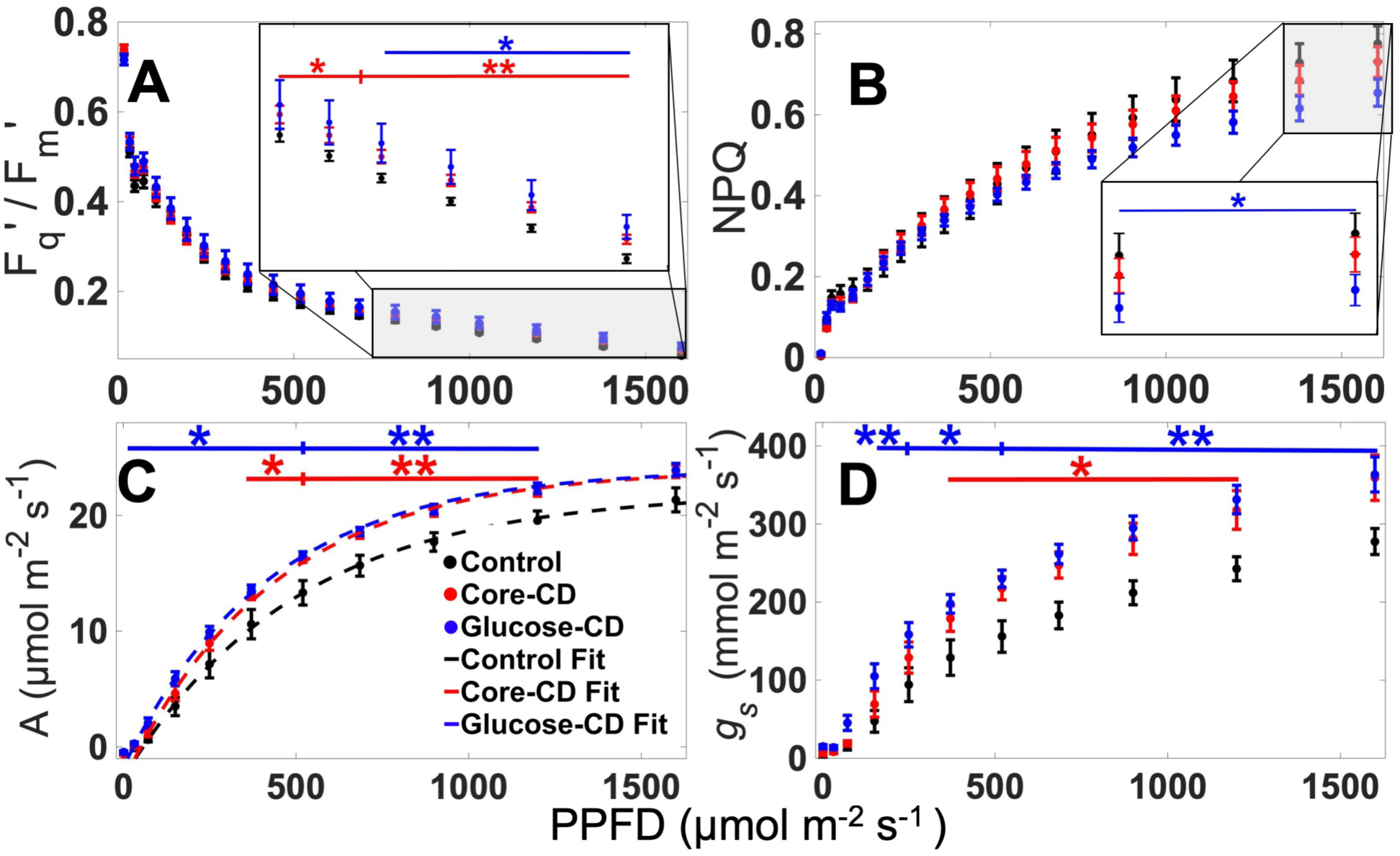
Irradiance dependent photosynthetic performance of *Triticum aestivum*. Light curves for (A) Photosystem II operating efficiency, (B) non-photochemical quenching, (C) carbon assimilation and (D) stomatal conductance obtained from chlorophyll fluorescence (A, B) and IRGA (C, D) measurements of the three different treatments. The light curve was fitted with an exponential model. The values of A_SAT_ are given in the Table 1. R^2^ ≥ 0.9945 for all fits.

The greater enhancement of carbon assimilation (*A*) compared to Photosystem II activity (*F*_*q*_’/*F*_*m*_’) observed for plants treated with CDs, indicates that they alter processes downstream from the initial light harvesting, such as electron transport chain and stomatal efficiency, to enhance photosynthesis.^16^ This hypothesis is supported by the increase in *A*_*SAT*_ upon addition of CDs, suggesting that the treated plants are able to perform more photochemistry with the light that they absorb, *e.g.* an increased operational efficiency, but do not enhance the maximum photosynthetic capacity of plants, since *A*_*MAX*_ is unaffected.

### Kinetics of photosynthetic performance

We then focused on the response of the treated plants to changes in light conditions by measuring the induction and relaxation of *A, g*_*s*_, *F*_*q*_’/*F*_*m*_’ and NPQ. These measurements were performed as dark-light-dark cycles as displayed in Figure 3. The induction of stomatal conductance was fitted using a previously developed model^17^ to quantify the maximum rate of stomatal conductance (Sl_MAX_) upon exposure to light. These experiments revealed that Sl_MAX_ increased for both of the CD treatments, Figure 3, B.

**Figure 3.**
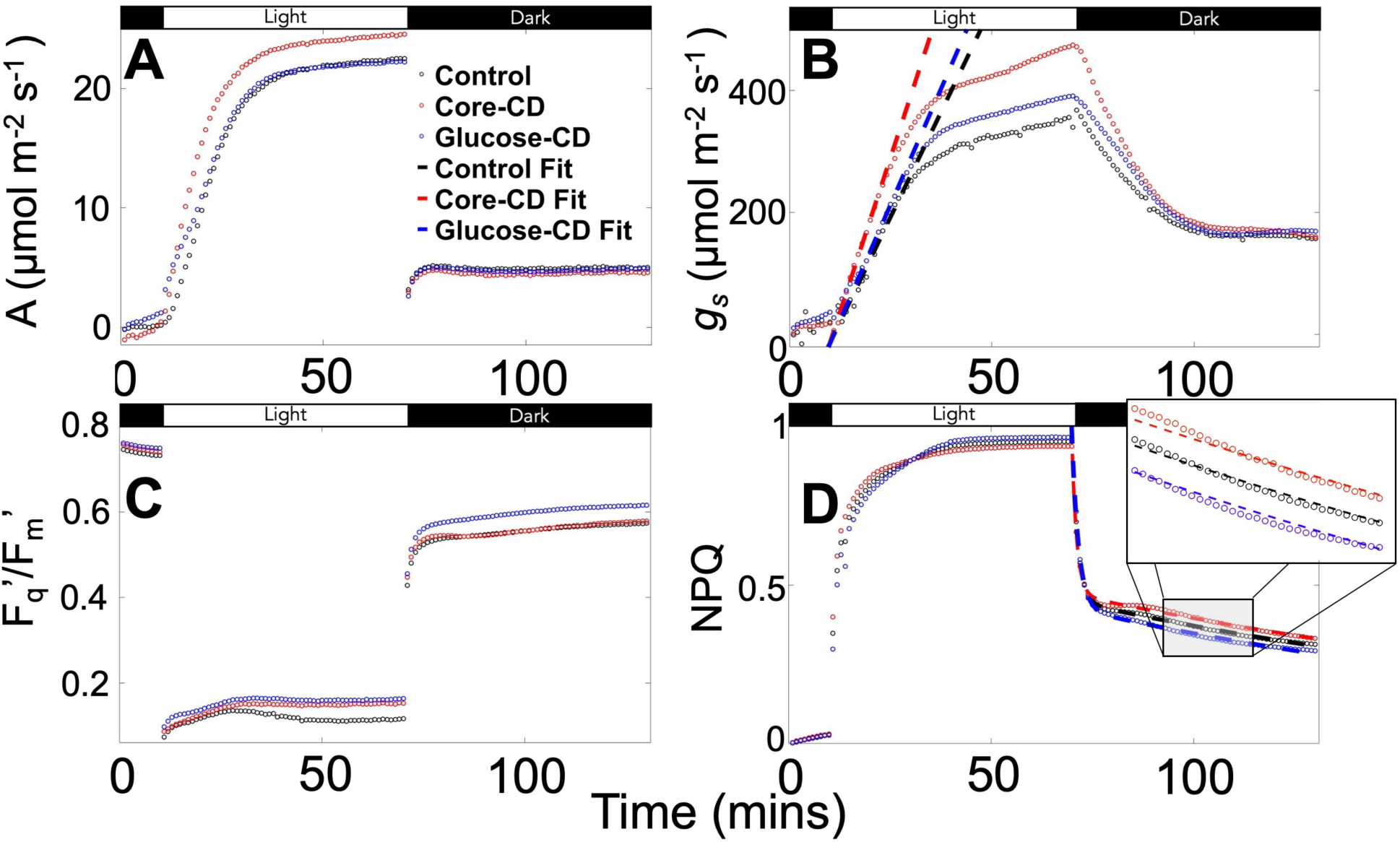
Kinetics of photosynthetic performance. Adaptation kinetics for (A) Carbon assimilation, (B) Stomatal conductance (Sl_MAX_ values were fitted using a previously developed model^16^), (C) Photosystem II operating efficiency, (D) NPQ kinetic where the relaxation of NPQ was fitted with a bi-exponential.^4^ The periods of dark and light are indicated with black and white boxes respectively. Light corresponds to a PPFD of 1000 μmol m^-2^ s^-1^ and dark corresponds to a PPFD of 0 μmol m^-2^ s^-1^ for (C, D) and 100 μmol m^-2^ s^-1^ for (A, B). Calculated values and details are given in the Table 1 and SI.

To quantify the relaxation of NPQ upon re-adaptation to dark conditions, the curves for 70-130 mins in Figure 3D were fitted to bi-exponential decays, as per previous studies,^4^ and tabulated in Table 1. Relative to untreated control, the rapid component of NPQ relaxation (NPQ_1_) was sped up by the core-CD treatment, yet glucose-CD treatment slowed down NPQ_1_. This is probably due to a relative reduction in the available carotenoid pigments which contribute to the fastest NPQ component, Figure 3, D (and confirmed by our pigment ratio analysis-see Figure 4).^4,18^ The relaxation of the secondary component (NPQ_2_), was slower (*e.g.* longer lifetime) for the core-CD treatment, implying a greater photoinhibition, which results in increased photodamage by the core-CD treatment, whereas NPQ_2_ is accelerated by the glucose-CD treatment, indicative of less photodamage - see Table 1.

**Figure 4.**
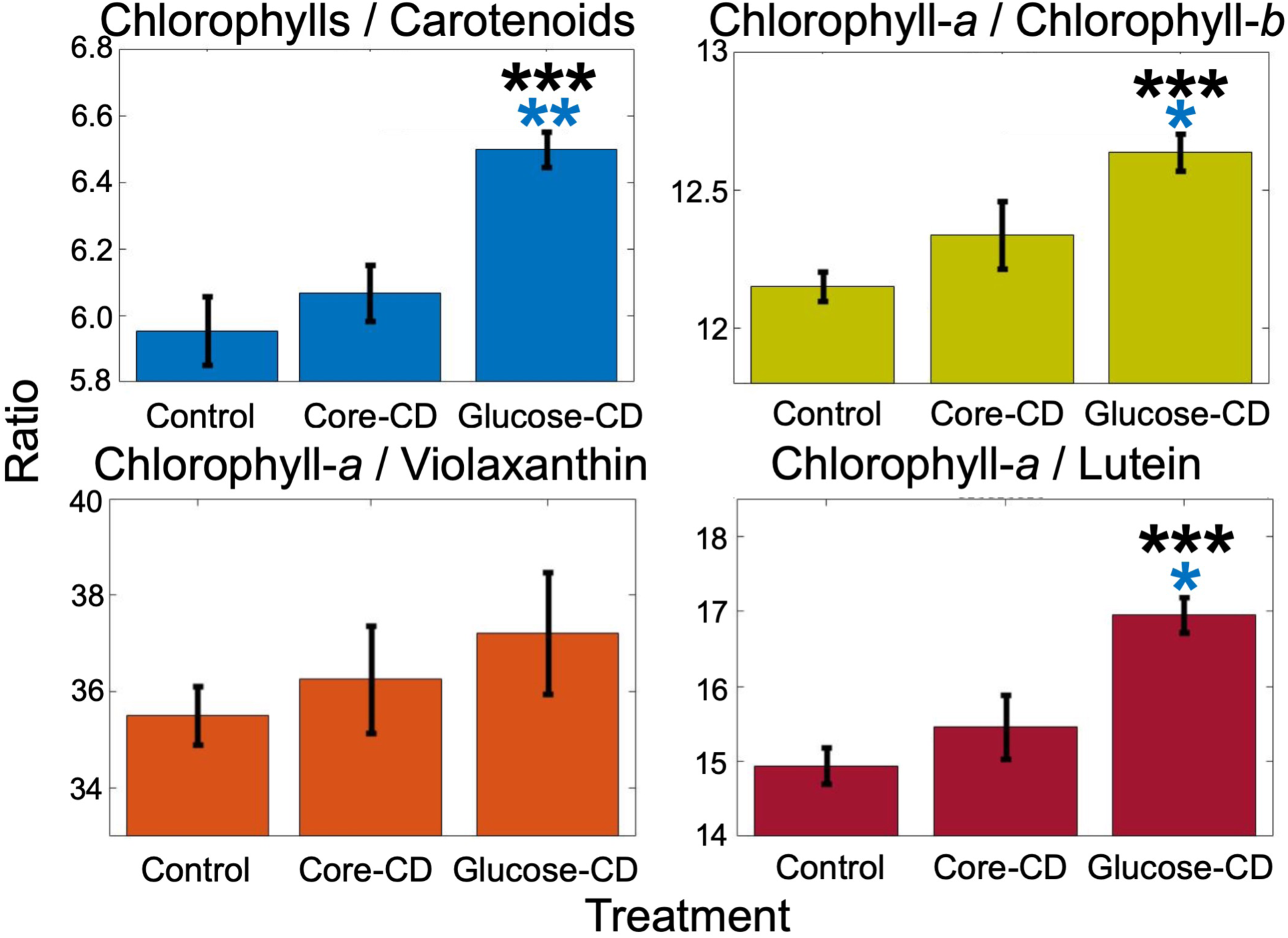
Photosynthetic pigment ratios. Pigment ratios extracted by HPLC from treated plants. Black asterisks indicate a difference from the untreated control, blue asterisks indicate a difference between the core and glucose CD treatments. Only the glucose-CDs demonstrated a significant difference from the control.

### Effect of CD treatment on Pigment Composition

The effect of CD treatment on the pigment composition was further investigated by the extraction and quantification of light harvesting and photoprotective pigments by high-performance liquid chromatography (HPLC), Figure 4. No significant differences in pigment ratios were observed between the untreated control and core-CD treatments. Conversely, for the glucose-CD treatment an increase in the ratio of chlorophylls to carotenoids, chlorophyll-*a* to chlorophyll-*b* and chlorophyll-*a* to lutein was observed compared to control and core-CD treatments. The increase in the ratio of chlorophyll-*a* to lutein supports the earlier conclusion that a reduced carotenoid pool in glucose-CDs treated plants slows the NPQ_1_ component of NPQ, although other processes that could affect NPQ_1_ cannot be completely disregarded.^19,20^ By contrast, no significant effect was observed on the ratio of chlorophyll-*a* to violaxanthin for either CD treatment. In general, the glucose-CD treatment results in an increased production of light harvesting pigments compared to photoprotective pigments, which suggests that glucose-CD treatment enables the plants to maintain reduced levels of photoprotection compared to the control or core-CD treatment.

The effects of this reduced photoprotection for the glucose-CD treatment were further investigated by measuring reactive oxygen species (ROS) production under illumination in isolated chloroplasts. Core-CDs were observed to increase ROS production compared to the untreated control (Figure S11), however, little effect was observed in the case of the glucose-CD treated plants, despite the glucose-CD treated plants producing fewer photoprotective pigments and performing less NPQ (see Figures 3,4). This suggests that the core-CD treatment results in increased photodamage compared to the control, which is prevented by the glucose-functionalization of the CDs.

### Changes to *Triticum aestivum* biomass production

The combination of effects on photosynthesis, photoprotection and photodamage leads to increased grain yield in *Triticum aestivum* treated with glucose-CDs, yet not the case for the core-CD treated plants. The total ear weight per plant for glucose-CD treatments are 18% greater than control (*p* < 0.01), the core-CD treatment was not significantly different from the control (Figure 5, B). The effect of the glucose-CD treatment was also demonstrated by an increase in seed production of 11% (*p* < 0.05) (Figure 5, C). A 17% increase in shoot biomass was also observed in glucose-CD treated plants (Figure S12). In all measurements, no significant effect was observed with treatment with glucose alone at the same concentrations (see SI).

**Figure 5.**
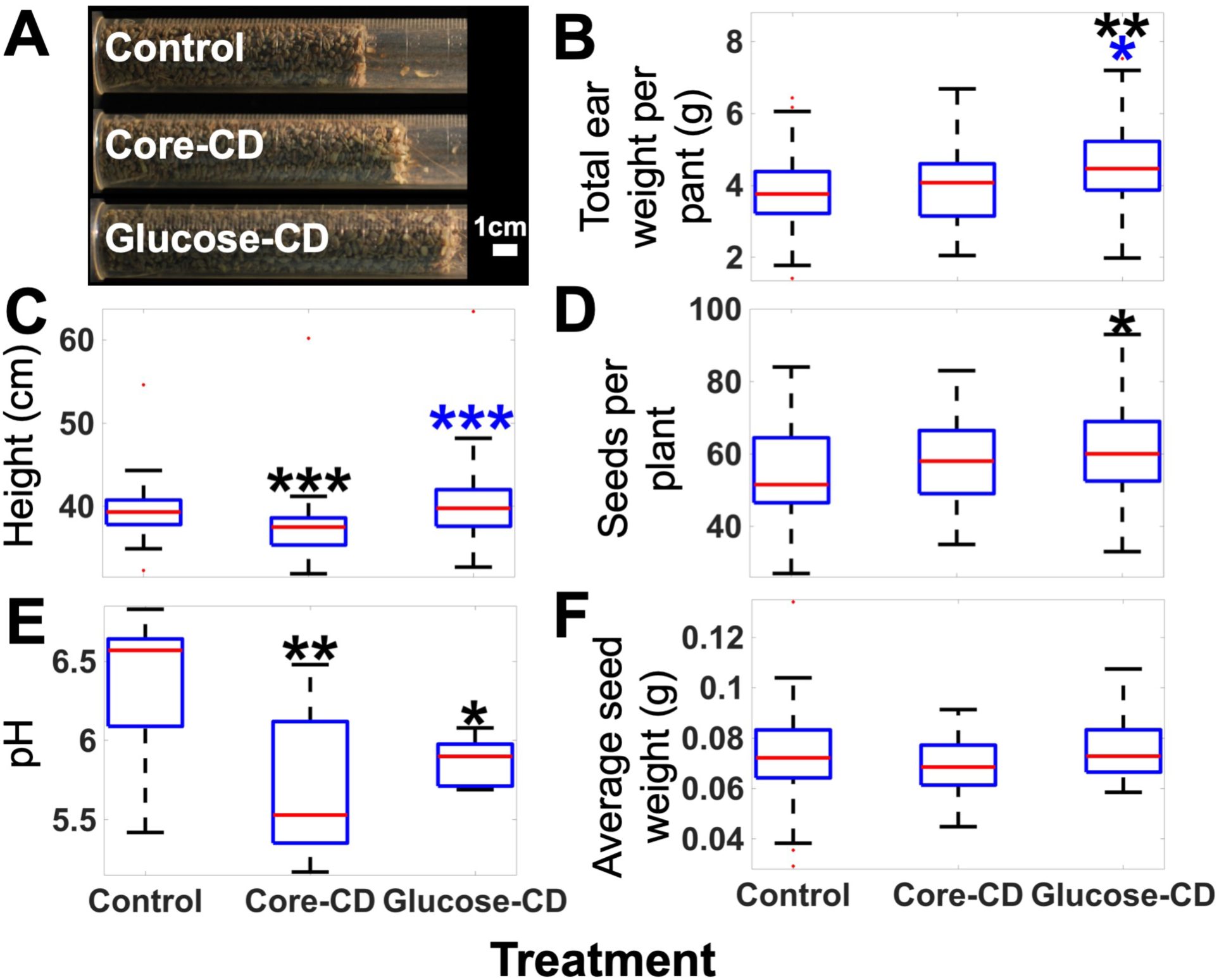
Physiology and yield measurements. (A) Image of the dried, threshed seed yield after treatments. For each box the red central line indicates the median, the blue top edges indicate the 25^th^ percentile, the blue bottom edge indicates the 75^th^ percentile, the whiskers indicate the range excluding outliers, red squares indicate outliers. Black asterisks indicate a difference from the control, blue asterisks indicate a difference between the core and glucose CD treatments.

## DISCUSSION AND CONCLUSIONS

Our results demonstrate that by utilizing biomolecule functionalized CDs it is possible to realize increased crop yields. While previously studies with CDs might have provided a route to the beneficial effect of enhancing photosynthesis, none have resulted in significant increased crop productivity: in agreement with what is observed with core-CD treatment in this study. This accentuates the need to design the functional surface of NPs for biological applications.

Furthermore, we explain the observed difference in crop yield between the core-CD and glucose-CD treatments are due to enhanced chlorophyll production; reduced NPQ by decreased lutein production; lowered photodamage demonstrated by reduced reactive oxygen species production; and enhanced uptake and internalisation into the mesophyll cells.

The methods demonstrated here provide a renewable, low-cost and facile route to an increase in *Triticum aestivum* productivity of 18% using functional nanomaterials. If we are to address the increase in crop productivity to meet forecast global food demand in the future,^21–24^ one potential route could be to embrace the use of nanomaterials to enhance yields. The use of designer-NPs has the potential to be combined with previously demonstrated transgenic techniques to bring crop productivity towards a theoretical maximum. These new techniques could also be applied to biofuels, biomass and bio-photovoltaics with the potential to enhance green-energy production.

We have demonstrated for the first time the use of glycan-functionalized nanomaterials to obtain an increase in the yield of crop. These materials also exhibit high bio-availability enabling simple treatment on a larger scale providing a significant advantage over previous methods. The synthetic routes used to access the nanomaterials are simple and sustainable. As a consequence, we believe this work represents a step towards the use of designer nanomaterials for agricultural applications.

## METHODS

### Statistics and significance

1, 2 or 3 asterisks are used throughout the manuscript to indicate a significant increase above the control of p ≤ 0.05, p ≤ 0.01 or p ≤ 0.001 respectively. An additional table of statistical parameters calculated from the physiological measurements is given in the SI (Table S2). For Figure 3 N=5 except for the control in (A, B) where N=4. For Figure 4 N=5. For Figure 5: (A) N=32, for (B, C and F) N=36, 32 and 36 for the control, core-CD and glucose-CD treatments, respectively; (D) N=35, 32 and 35 for the control, core-CD and glucose-CD treatments, respectively and (E) N=9.

### CD synthesis and characterization

Core CDs were synthesized following a modified procedure from our group.^11,14^ The glucose functionalized CDs were prepared via a two-step procedure starting from core-CDs. In brief, treatment of core-CDs with succinic anhydride to generate acid-coated CDs, followed by amide conjugation with 1-amino glucose gave after purification through a 200 nm syringe filter and size-exclusion chromatography (Sephadex G-10, Sigma Aldrich) the functionalized CDs (See ESI for full experimental details). Glycan conjugation was performed with an excess of 1-amino glucose to ensure all the acid groups reacted. This solution was then stirred vigorously overnight. For storage, the glucose-CDs were dissolved in methanol and kept at 4°C to prevent aggregation. See the SI; scheme S1 and Figure S1-S6 for full details of the synthesis; NMR; Fourier transformed infrared spectroscopy; dynamic light scattering; absorbance spectroscopy and fluorescence spectroscopy.

### Plant growth conditions and treatments

The plants were grown in compost (Levington, F2) in a greenhouse at a constant 20.0 °C temperature. Natural lighting was supplemented with LED lighting (PhytoLux, Attis-7) 80–120 μmol m^2^ s^-1^ to provide a 16-hour photoperiod. From three weeks post germination until harvesting the plants were treated three times a week with 3.3 mg of treatment per plant, totalling 10 mg of treatment per plant per week. Productivity measurements were made 10 weeks post germination at Zadoks stage 9.0–9.2 and after 7 weeks of treatment.

### Fluorescence microscopy

Samples were imaged with a Leica DM200 and Leica MC120HD detector. Chlorophyll fluorescence was imaged using 470 nm excitation and collection of emission between 650**–**700 nm, the CDs were imaged with a 365 nm excitation and 430**–**470 nm emission. Leaf tissues were imaged 7 weeks post-germination and after 4 weeks of treatment. To demonstrate the CD fluorescence each treatment was imaged using the same excitation, microscope and detector settings. Histograms of the fluorescence observed across three images for each treatment across are given in Figure S8.

### Chlorophyll fluorescence and gas exchange measurements

A GFS-3000 (Waltz) and a MAXI-PAM (Waltz) were used for IRGA and chlorophyll fluorescence respectively. Measurements were made at Zadoks 7.3–7.7 on flag leaves at seven weeks post germination. Plants were dark adapted for 40 minutes before light curve measurements. For chlorophyll fluorescence, the average across 10 equally sized areas evenly distributed across the leaf was recorded for each value. For the light curves the actinic light step length was 30 seconds. [CO_2_] of 400 μmol mol^-1^, [H_2_O] of 17 mmol mol^-1^, leaf temperature of 22 °C and vapour pressure deficit of 1.0 ± 0.2 kPa were all maintained for IRGA experiments.

### Pigment measurements

Pigments were extracted from flag leaves and analysed by HPLC using previously developed methods.^25^ All solvents used were HPLC-grade. Samples were taken at the middle of the photoperiod on the same day, 0.1g of flag leaf was used for each sample, *N*=5. A stationary phase 3.5 µm spherical silica particle with an 80 Å pore size was used (Agilent, Eclipse XDB C8, 4.6 mm × 150 mm). 430 nm, 470 nm and 450 nm were used to detect the absorbance of chlorophyll-*a*, chlorophyll-*b* and the carotenoids respectively.

Due to the dark adaption of the leaves no zeaxanthin or antheraxanthin were observed in the samples. It was not possible to separate α and β carotene, so peaks are simply labelled carotene however is expected to be predominantly β-carotene.

## DATA AND SOFTWARE AVAILABILITY

The accession number for the [data or structure] reported in this paper is [database abbreviation]: [accession number].

## SUPPLEMENTAL INFORMATION

Supplemental Information includes twelve figures, one scheme and two tables.

## ACKNOWLEDGMENTS

We would like to thank Tom Pitman, Alanna Kelly and Anna Lim (University of Bristol) for growing the plants; Taryn Fletcher (University of Bristol) for her help with experimental work; and James Stevens (University of Essex) for his advice on IRGA measurements. This research was supported by EPSRC-BCFN PhD studentship EP/G036780/1 and EPSRC Doctoral Prize fellowship EP/R513179/1 (TAS); EPSRC CAF EP/J002542/1 and ERC-COG:648239 (MCG); NERC grant NE/N006518/1 (MLY, DF); BBSRC grants BB/NO16831/1 & BB/N021061/1 (TL), Royal Society University Research Fellowship UF1402310 (TAAO), the Bristol Centre for Agricultural Innovation and the Wolfson Foundation.

## AUTHOR CONTRIBUTIONS

TAS, TAAO, MCG, TL and HMW conceived the experiments; TAS carried out the microscopy, chlorophyll fluorescence, CD-pigment interactions, IRGA and physiology experiments; TAS and DF performed the HPLC; DBA developed the modified synthesis; SAH created the original synthesis; TAS synthesized and characterized the CDs; and TAS, TAAO, HMW and MCG co-wrote the manuscript, which all authors commented on and edited.

## DECLARATION OF INTERESTS

The concepts of the paper are shared with a patent filed by the University of Bristol: International (WO) Patent Application No: PCT/EP2018/097143 on which TAS, DBA, MCG and HMW are named inventors.

