## supplementary information for "Photosynthesis and crop productivity is enhanced by glucose-functionalized fluorescent carbon dots"

### 1 General

NMR figures were made using MestReNova, all other plots were produced using MatLab. Schemes, chemical structures and synthesis were produced in ChemDraw. Analysis of microscopy was performed using ImageJ. All fluorescence measurements were recorded using a Perkin-Elmer LS45 spectrometer in a 3mm path length quartz cell (Thor Labs.). All quoted values are means. For scatter or column plots error bars shown correspond to the standard error. When comparing the glucose treatment to the control 2-tailed tests were used. When comparing the CD treatments to the control 1-tailed tests were used. When comparing the core-CD and glucose-CD treatments 2-tailed tests were used. For all measurements either N<15 or distributions were observed to be near Gaussian therefore unpaired t-tests were used to calculate p-values. The adjusted R-squared statistic ($\overline{R^2}$) was used to determine the quality of fitting. For all fitting, the non-linear least squares method was used with a trust-region algorithm. When fitting, all treatments were given the same start points and boundaries. 1, 2 or 3 asterisks are used to indicate a significant increase above the standard control of $p \leq 0.05$, $p \leq 0.01$ or $p \leq 0.001$ respectively, the colour is used to indicate the treatment being compared to the control.

### 2 Synthesis

For the synthesis chemicals were purchased and used without further purification and samples were concentrated under reduced pressure using both a rotary evaporator (Büchi) at a pressure of 15 mmHg (diaphragm pump) at room temperature. Where possible HPLC-grade solvents were used.

The original synthetic route for these CDs were originally developed by Hill *et al.* [15] this has since been modified by Swift *et al.*[12] and is again improved here. Details of differences from these previous synthetic routes are given below.

A simplified scheme of the synthesis of the CDs as well as a full description of the modified glycan functionalisation are given below.

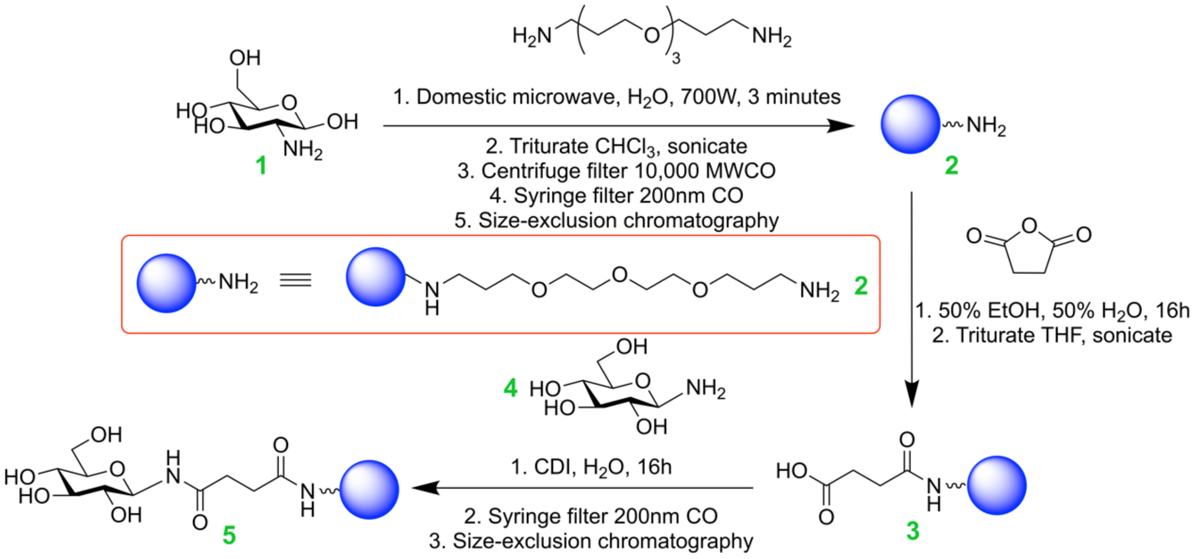

***Scheme 1:*** *Synthesis of the CDs. Each product is assigned a number, shown in green.*

To form the glucose functionalized CDs (**5**), 1-amino glucose (**4**) is bonded to acid decorated CDs (**3**), which have been synthesised using the previously described method,[11] by an amide bond. To achieve this the previous solution of acid decorated CDs (**3**) was passed through a 200nm syringe filter and mixed with 2 equivalents by weight of N-(3-Dimethylaminopropyl)-N'-ethylcarbodiimide (EDC) and 10 equivalents by weight of 1-amino glucose (**4**). Glycan conjugation was performed with an excess of 1-amino glucose to ensure all the acid groups reacted. This solution was then stirred vigorously overnight. The sample was then passed through a 200 nm syringe filter and. This was then purified by size-exclusion chromatography (Sephadex G-10, Sigma.) The CDs were identified as a brown band on the column and were identified by both fluorescence and absorbance spectroscopy, both tails of the CD band were discarded, particularly the lower molecular weight tail as this may contain non-functionalized CDs. The resulting fraction was freeze-dried, weighed and suspended in methanol.

For storage, the glucose-CDs (**5**) were dissolved in methanol and kept at 4˚C to prevent aggregation.

### 3 NMR spectroscopy

The $^{1}H$ and HSQC 500MHz NMR spectra of the CDs are shown below in figures S1-S2. All samples were dissolved at 5 mg/mL in 0.8 mL of $D_2O$ using Norrell Select Series 7" NMR tubes (S-5-500-7). All spectra were taken on a Bruker Advance III HD 500 Cryo. All shifts are quoted in ppm. Residual internal $D_2O$ is at $\delta$ = 4.70. Peaks were identified using both $^{1}H$ and HSQC NMR. Coupling constants (J) given in Hertz. Multiplicities are abbreviated as: s (singlet), d (doublet), t (triplet), q (quartet), p (pentet) and m (multiplet).

The $^{13}C$ and $^{1}H$ NMR spectra of the core CDs shown in Figure S1 and reproduce our previously reported data.[10,11] The peaks arising from the TTDDA linker are identified and assigned. C=C bending and stretching, Aryl C-O stretching and a weak phenol O-H bending signal can also be observed in the FTIR to support this conclusion.

The NMR spectra for the glycan-functionalized CDs are shown in figures S2-S6. For each of these spectra the TTDDA linker is identified, although often not completely assigned due to overlapping peaks from the functionalized carbohydrates (hydrogens attached to carbons 2-6). These peaks are themselves also difficult to assign individually due to their overlap, however appear in the following characteristic ranges: $^{1}H$ $\delta$ = 3.0-3.8 and C-13 $\delta$ = 55-80. Due to the apparent congestion in the NMR spectra, the successful carbohydrate functionalisation was instead determined by the identification of the characteristic doublet peak associated with the hydrogen on the anomeric carbon of intact carbohydrates and which is shifted when compared to the unconjugated glycoside. This doublet is found in the following ranges: $^{1}H$ $\delta$ = 4.2-5.5 and $^{13}C$ $\delta$ = 90-105. Often more than one doublet is observed, which arises from reduced conformational freedom that the carbohydrate experiences due to being tethered to the nanoparticle surface. This agrees with our previous NMR studies.[11,12]

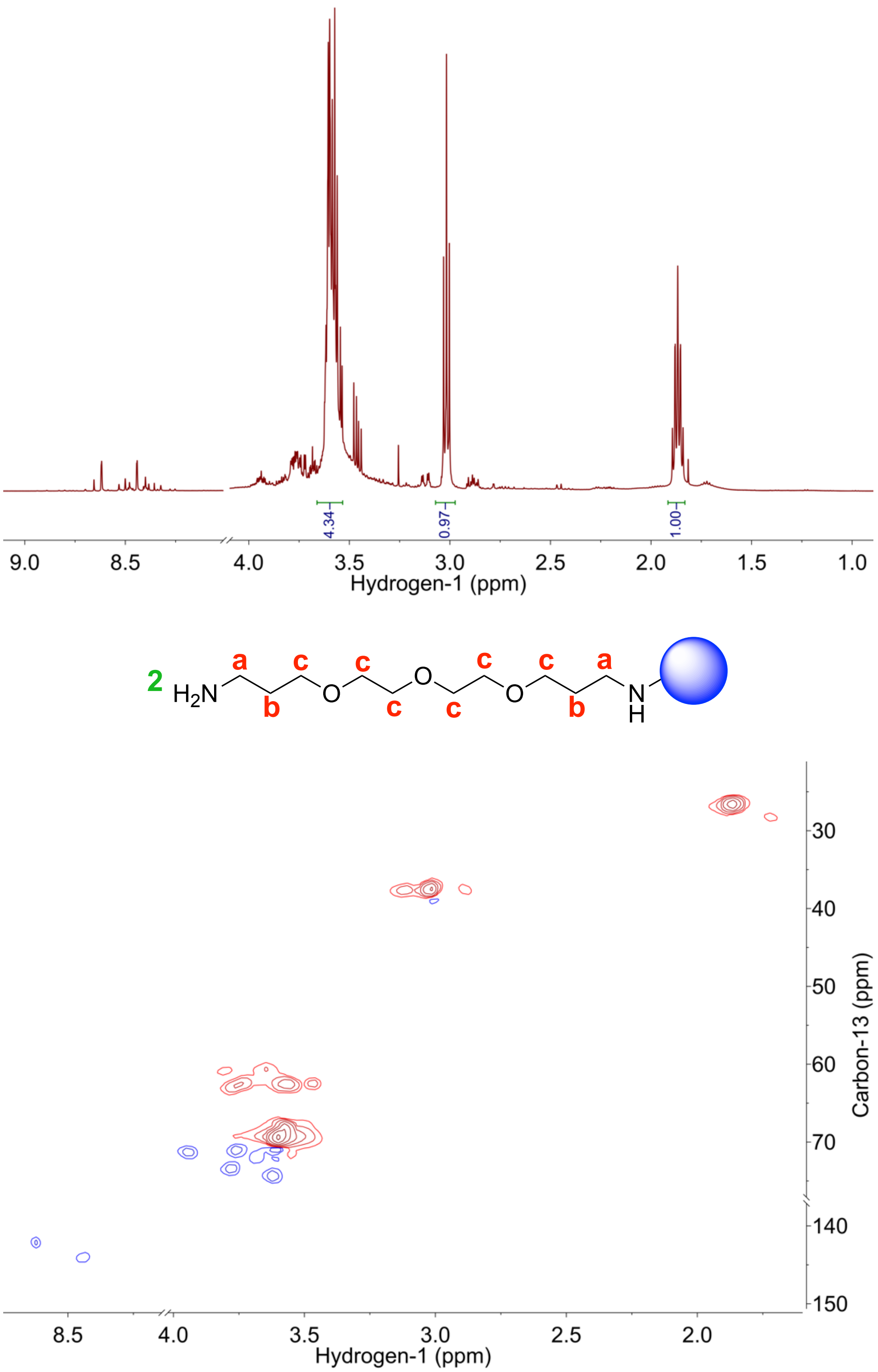

***Figure S1:*** *$^1$H (top) and HSQC (bottom) NMR spectra of the core-CDs. $\delta$: 3.58 (m, 16H, H-c), 3.02 (t, J = 7.2 Hz, 4H, H-a), 1.87 (q, J=6.0, 4H, H-b).*

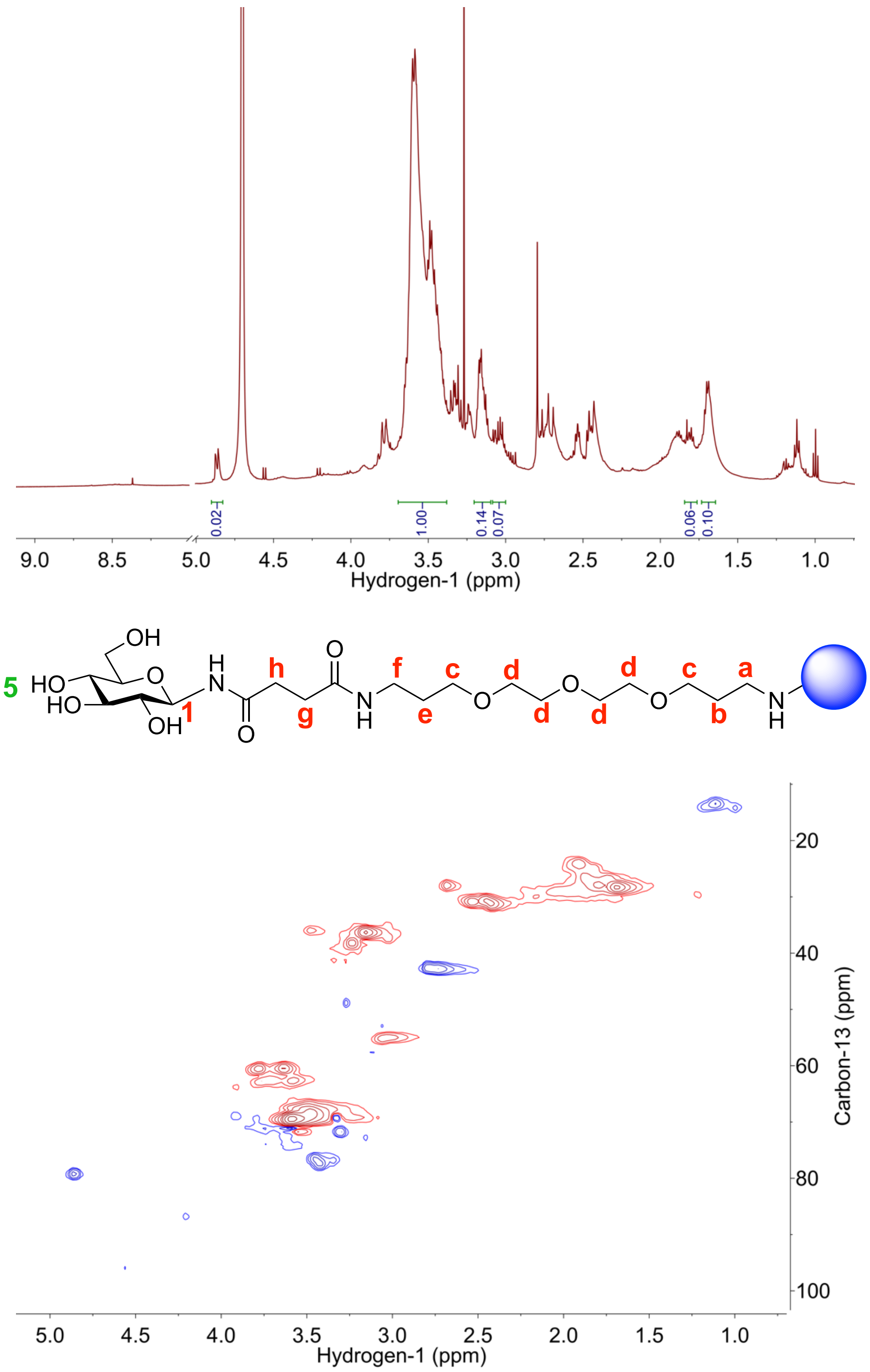

***Figure S2:** $^{1}$H (top) and HSQC (bottom) NMR spectra of the glucose-CDs. $\delta$: 4.86 (d, 1H, H-1), 3.32 (t, 12H, H-c, and H-d), 3.14 (t, 2H, H-f), 3.03 (t, 2H, H-a), 1.80 (t, 2H, H-b), 1.63 (p, 2H, H-e).*

### 4 Fourier transformed infrared spectroscopy (FTIR)

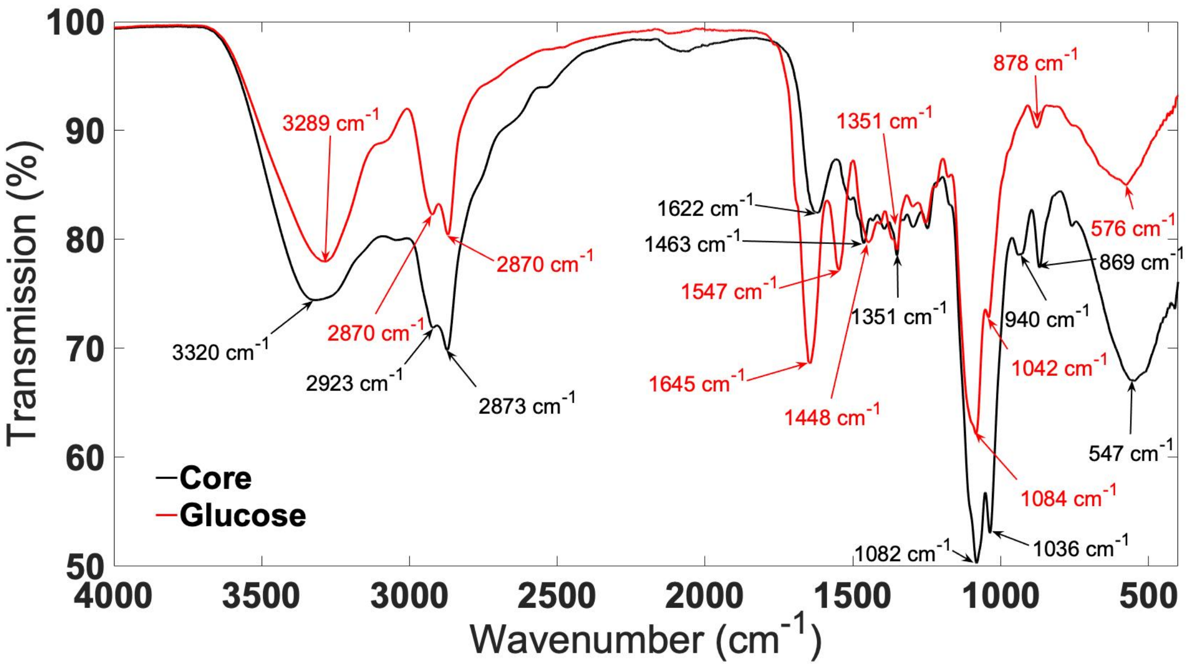

***Figure S3:*** *The following peaks are identified Core-CDs: 547cm$^{-1}$ (C-Cl stretching); 869cm$^{-1}$ and 940cm$^{-1}$ (C=C bending); 1036cm$^{-1}$ (ether C-O stretching); 1082cm$^{-1}$ (primary and secondary alcohol C-O stretching); 1351cm$^{-1}$ and 1463cm$^{-1}$ (alcohol O-H bending); 1622cm$^{-1}$ (amide C=O stretching); 2873cm$^{-1}$ and 2923cm$^{-1}$ (sp$^3$ C-H stretching); 3320cm$^{-1}$ (alcohol O-H or amine N-H stretching). Glucose-CDs: peaks identified as: 576cm$^{-1}$ (C-Cl stretching); 878cm$^{-1}$ (C=C bending); 1042cm$^{-1}$ (ether C-O stretching); 1084cm$^{-}$ (primary and secondary alcohol C-O stretching); 1351cm$^{-1}$ and 1463cm$^{-1}$ (alcohol O-H bending); 1547cm$^{-1}$ (C=C stretching); 1645cm$^{-1}$ (amide C=O stretching); 2873cm$^{-1}$ and 2923cm$^{-1}$ (sp$^3$ C-H stretching); 3320cm$^{-1}$ (alcohol O-H or amine N-H stretching). Spectra were recorded on a Bruker ATR.*

### 5 Absorbance Spectroscopy

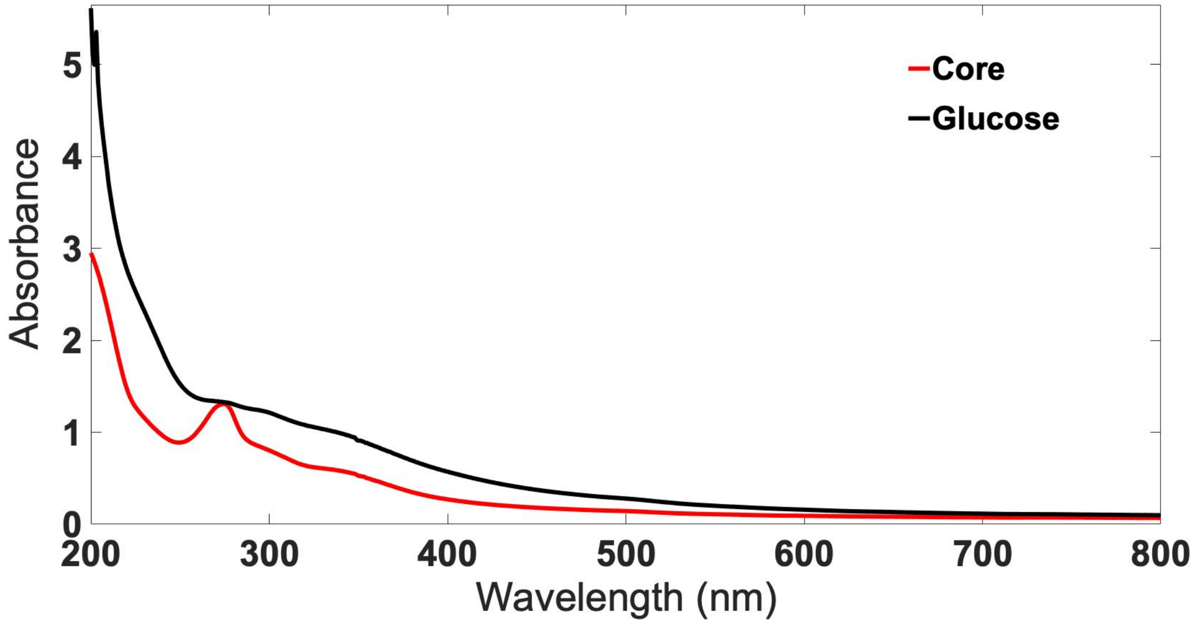

***Figure S4:*** *UV-visible absorption spectra for the CDs, spectra were recorded at a concentration of 1.0 mg/ml in methanol. The spectra were recorded using a Cary UV-Visible 50 spectrophotometer and are dominated by bands at 273 and ≤200 nm.*

### 6 Fluorescence Spectroscopy

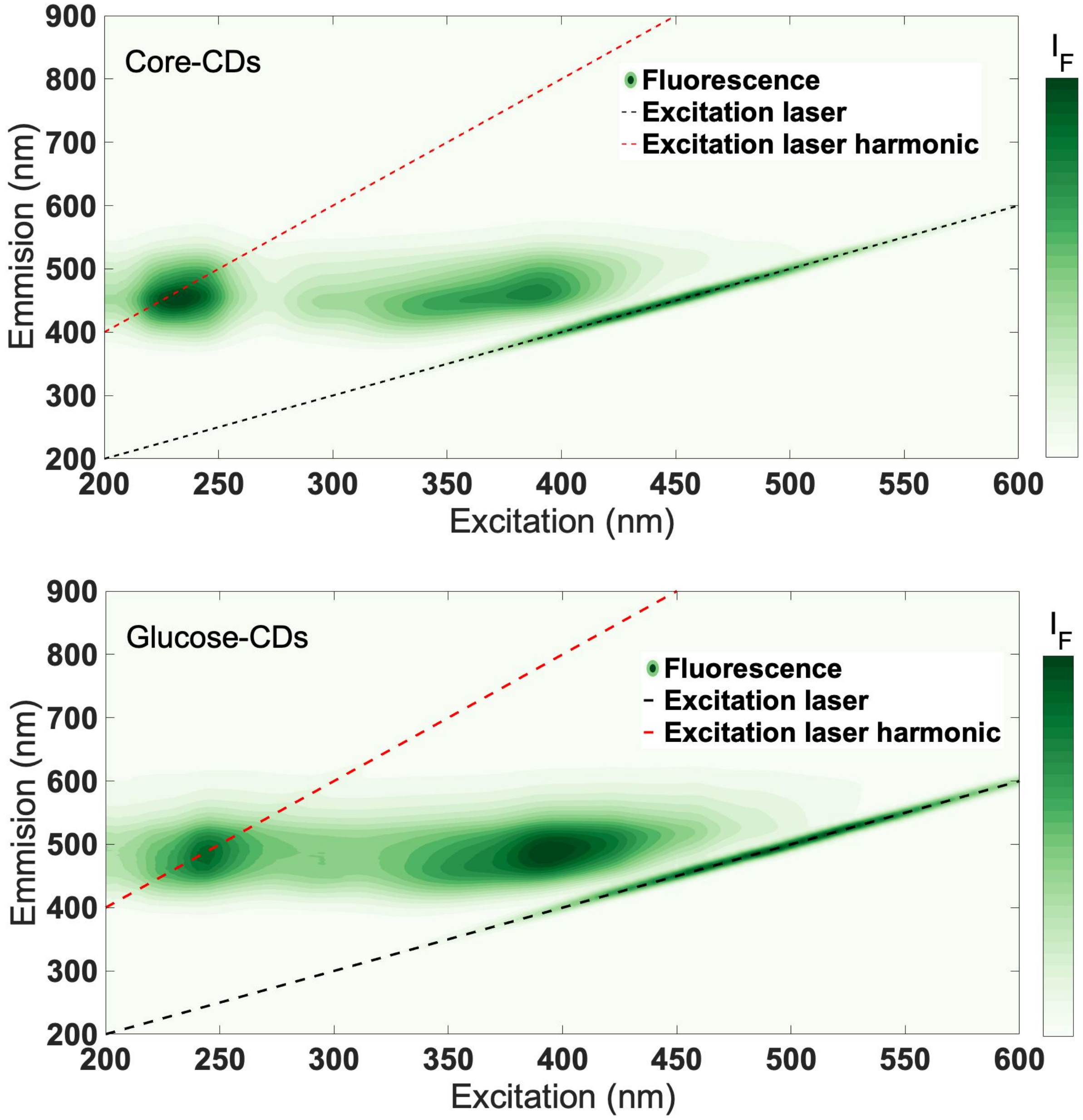

***Figure S5:*** *Two-dimensional excitation-emission correlation fluorescence spectra of the Core-CDs (top) and the Glucose-CDs (bottom). Spectra were acquired at 1.0 mg/ml CD in water. A full discussion of the fluorescence properties is given by Swift et al.*[11]

### 7 Dynamic Light Scattering (DLS)

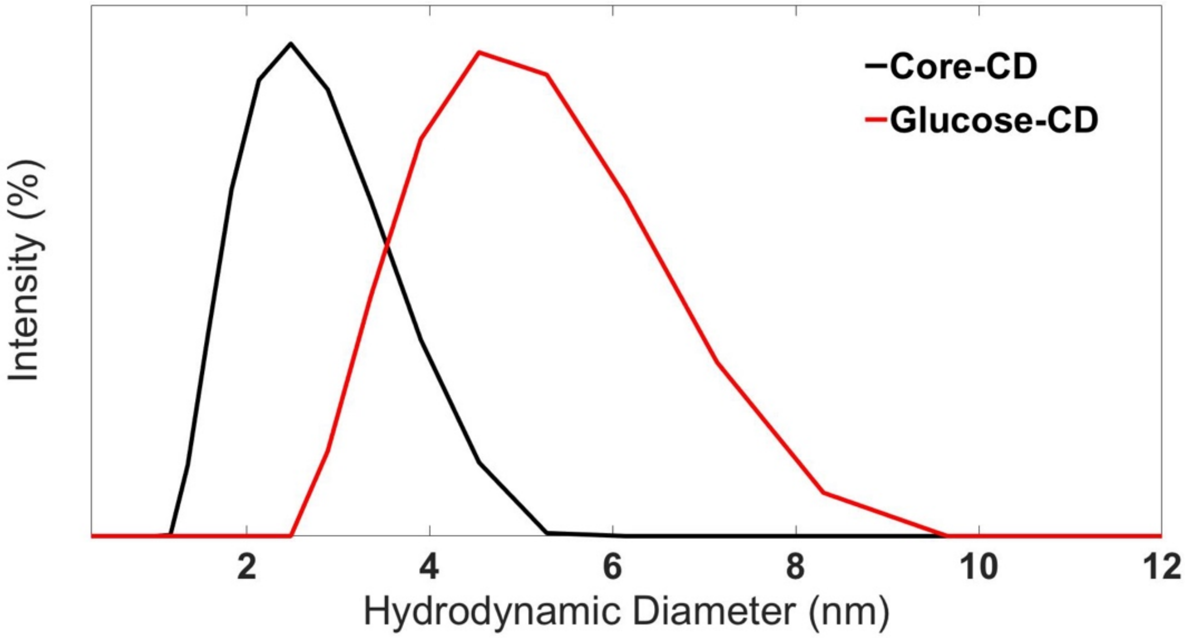

***Figure S6:*** *DLS spectra for the CDs in HPLC-grade methanol at a concentration of 1.0 mg $ml^{-1}$. Data was collected using a Malvern Zetasizer Ultra.*

### 8 Fluorescence Microscopy

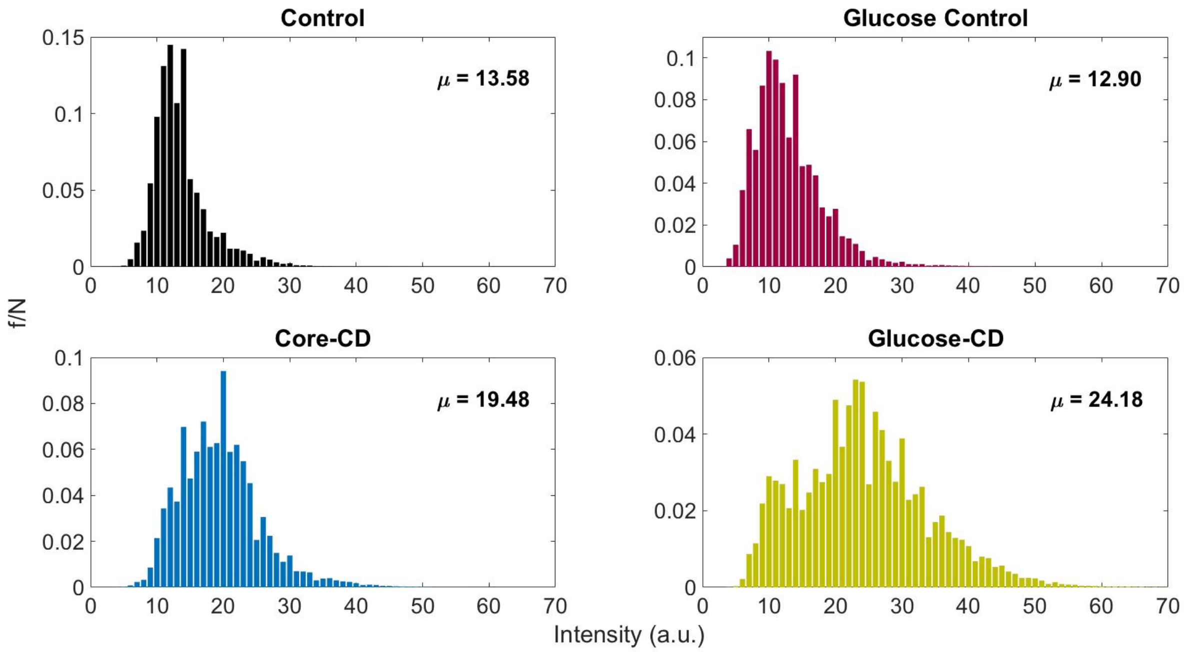

***Figure S7:*** *Histograms of the fluorescence observed for each treatment. N=7,485,696 pixels across three images, each taken from separate leaves of similar thickness, the mean is quoted on each histogram.*

### 9 Quantification of CD Uptake

The uptake of the CDs was investigated by re-extraction of the CDs from leaf tissue and quantification by fluorescence spectroscopy and comparison with a calibration.

2g of leaf tissue was collected, flash frozen with liquid nitrogen and stored at -80°C. For analysis the tissue was thoroughly homogenised, suspended in HPLC-grade water and put into dialysis bags (10,000 MWCO) and dialysed against HPLC-grade water for 3 days. The contents of the dialysis bags were discarded and dialysis liquid was reduced *in vacuo*. The resulting solution was then separated by size-exclusion chromatography (G10 sephadex), the CD containing band was identified visually as a brown band and confirmed by fluorescence and absorbance spectroscopy, the entirety of this fraction was then freeze-dried and resuspended in 0.5ml of HPLC grade methanol.

The fluorescence from those samples was then compared with a calibration plot measuring fluorescence of each CD vs concentration in μg/mL (Figure S8). In addition, to correct for extraction/manipulation losses that can lead to underestimation of CD concentration in the leaves, the values were corrected by an extraction efficiency coefficient (EEC). This effect was quantified by adding a known amount of CDs with homogenised untreated leaf tissue at a concentration of 25μg of CD per gram of leaf tissue. These CDs were then extracted from the leaf tissue using the same methods, giving yields for the extraction process of 81.2% and 80.7% respectively, N=5.

Uncorrected values gave leaf concentrations of 23±1μg Core-CD per gram of leaf tissue and 26±1μg Glucose-CD per gram of leaf tissue, N=15. The Glucose-CD concentration was observed to be greater than the Core-CD concentration, $p=6.27x10^{-7}$. After the ECC correction is applied, the adjusted leaf concentrations are 29±2μg Core-CD per gram of leaf tissue and 32±1μg Glucose-CD per gram of leaf tissue. The Glucose-CD concentration was observed to still be greater than the Core-CD concentration, $p=5.47x10^{-7}$.

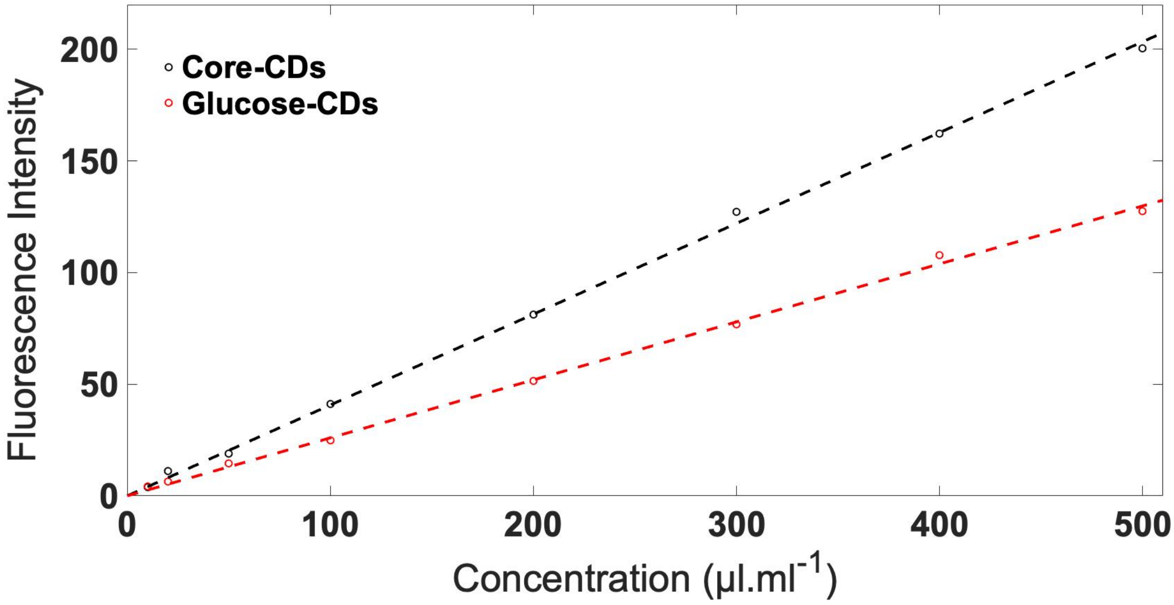

***Figure S8:*** *[CD]-fluorescence emission calibration plot. Fluorescence was detected at 445nm from 340nm excitation and averaged for 60 seconds for each measurement.*

### 10 Infrared Gas Exchange (IRGA)

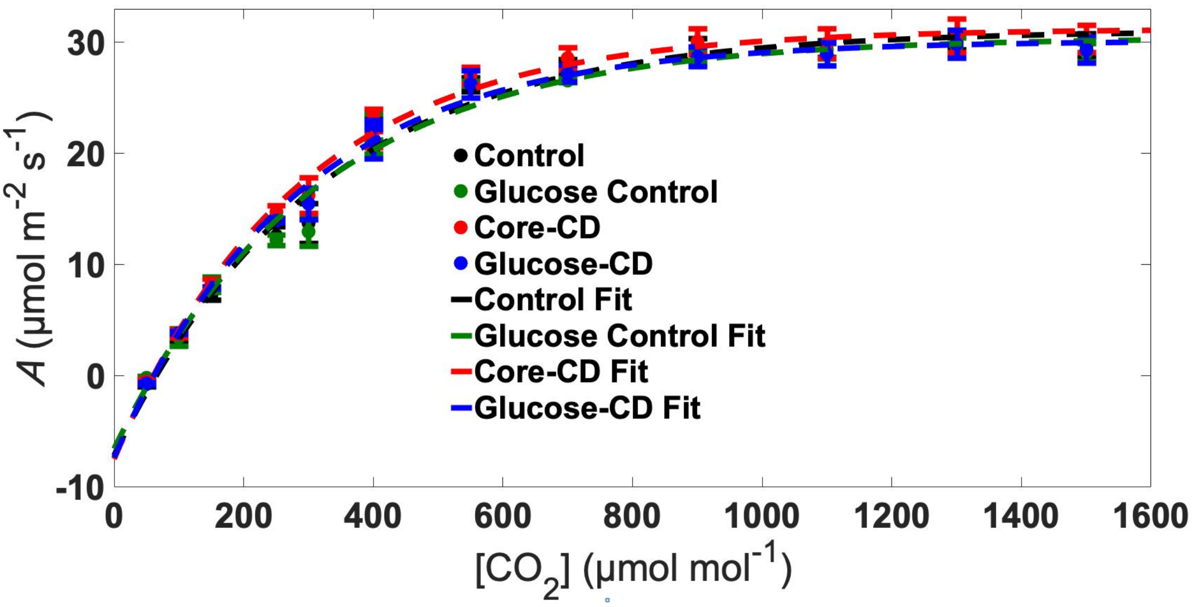

***Figure S9:*** *A-[$CO_2$] curve of all treatments, N=5.*

### 11 Extraction of pigments and analysis by HPLC

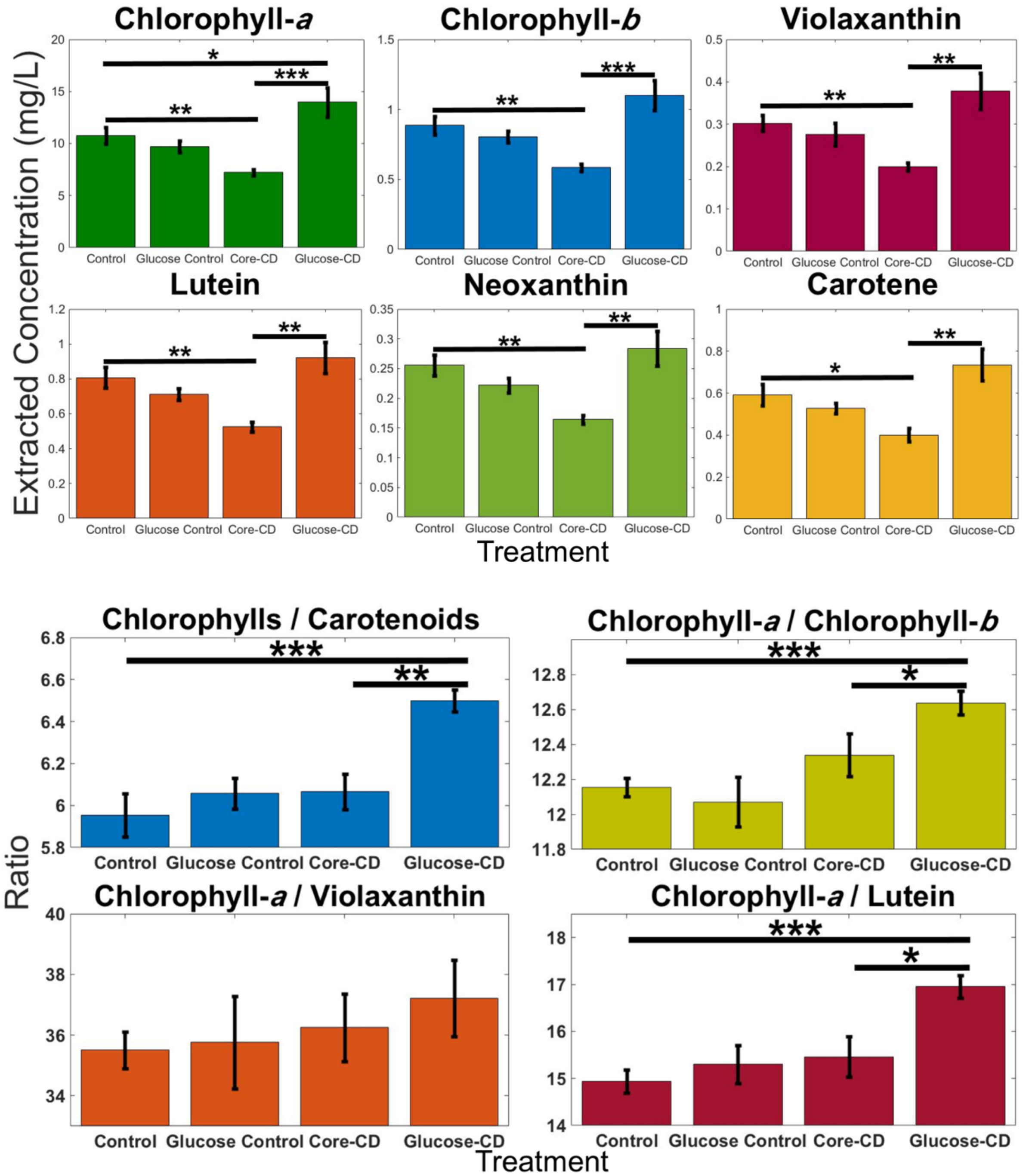

***Figure S10:*** *Extracted concentrations of all observed pigments and pigment ratios of the CD-treated Triticum aestivum, N=5.*

#### 12 ROS production in extracted chloroplasts

To investigate the interactions between CDs and ROS production chloroplasts were extracted from plant material using the same methods as highlighted in section 9 and based on those utilised by Giraldo *et al.*[6] CellROX Orange (Thermo Fisher) was used as a ROS sensitive dye with a peak fluorescence excitation at 545nm and emission 565nm.

Chloroplasts at 10$\mu$gml$^{-1}$ and CDs at various concentrations were incubated with 5 μM of CellROX Orange. Chloroplasts were kept in250 $\mu$molm$^{-2}$s$^{-1}$ PPFD. Controls of just the core-CDs, glucose-CDs, and chloroplasts as well as positive (Tert-butyl hydroperoxide) and negative (N-acetylcysteine) control were included.

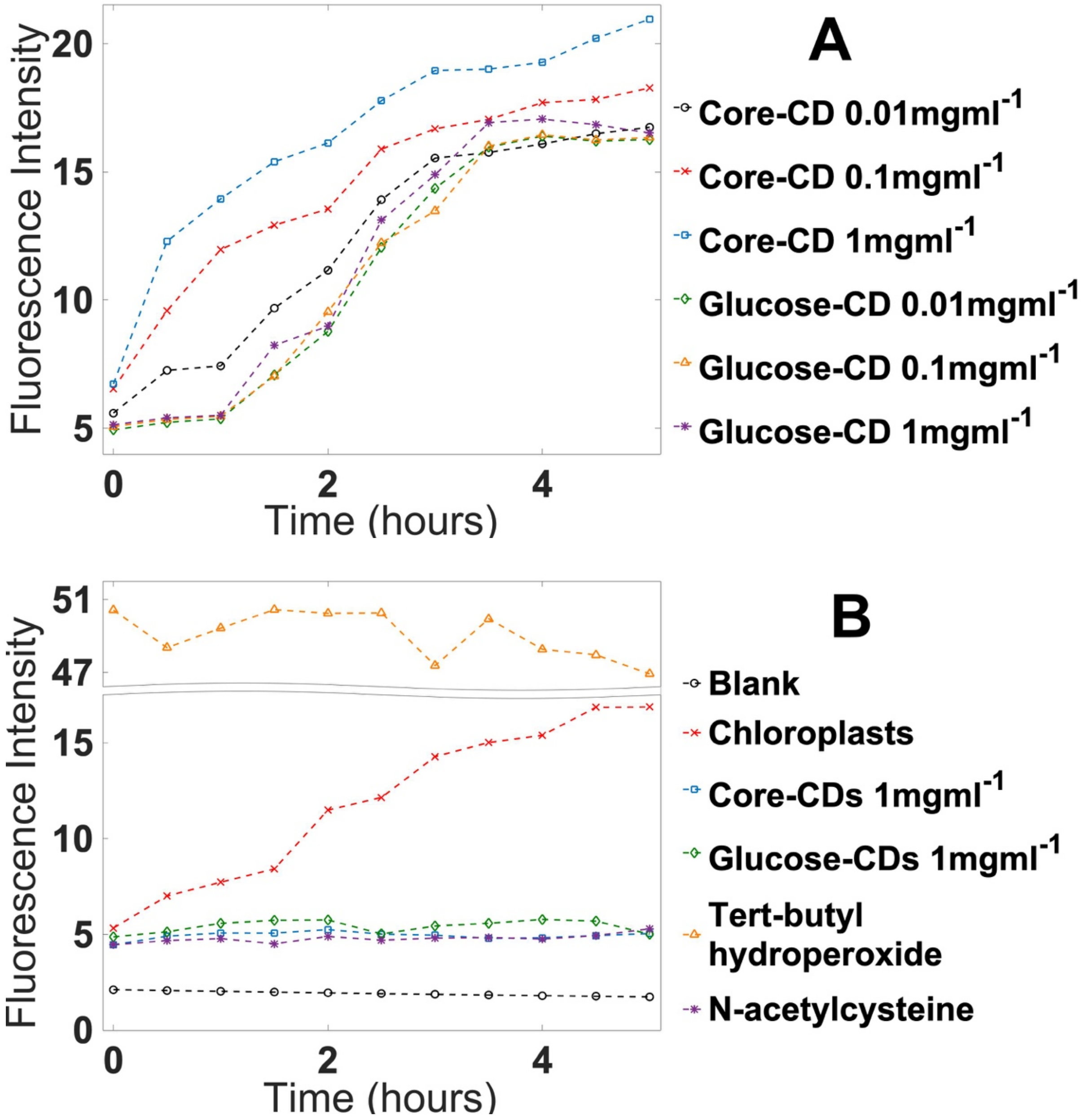

***Figure S11:*** *(**A**) CellROX fluorescence as a result of ROS production in chloroplasts due to different CD treatments. (**B**) CellROX fluorescence as a result of ROS production in a series of controls including untreated chloroplasts, CDs without chloroplasts and a blank cuvette.*

#### 13 Physiology and pH

To measure the pH of the soil 5g of soil was mixed vigorously with 25ml of water (MilliQ double distilled, pH=7). This mixture was left to stand for 30 minutes then mixed vigorously. The pH was then recorded using a calibrated pH probe (Mettler Toledo, SevenEasy). Statistical values for the physiology are included in Table S2.

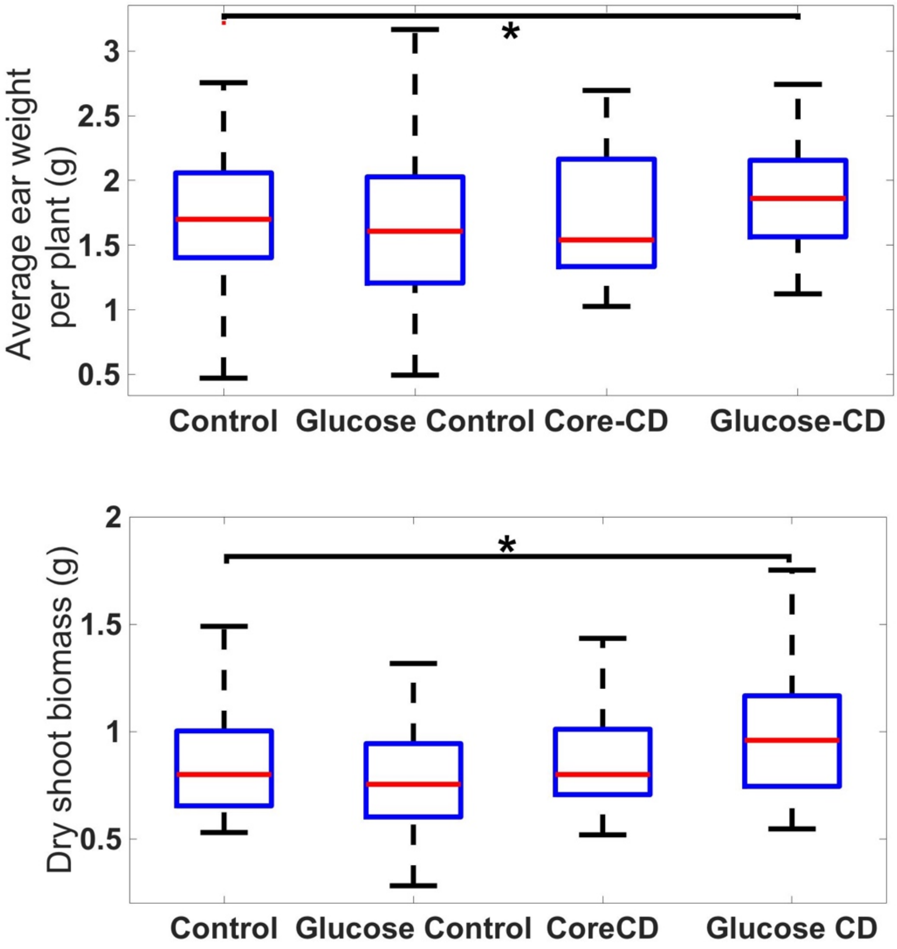

***Figure S12:*** *Average dry shoot biomass and average ear weight for each of the treatments. Biomass was dried in a 90℃ oven for 11 days.*

| Value | | Treatment | | | |
|---|---|---|---|---|---|
| | | Control | Glucose Control | Core Carbon Dot | Glucose Carbon Dot |
| Dry Shoot Biomass (g) | $\mu$ | 0.83 | 0.79 | 0.86 | 0.97 |
| | SD | 0.23 | 0.24 | 0.24 | 0.27 |
| | SE | 0.04 | 0.04 | 0.04 | 0.05 |
| | p value | | 0.545 | 0.520 | 0.0198 |
| | Percentage | | -4.1 | 4.4 | 17.1 |
| | N | 35 | 36 | 33 | 36 |
| Soil pH | $\mu$ | 6.3 | 6.3 | 5.7 | 5.86 |
| | SD | 0.5 | 0.3 | 0.5 | 0.15 |
| | SE | 0.2 | 0.1 | 0.2 | 0.05 |
| | p value | | 0.853 | 0.00317 | 0.0249 |
| | Percentage | | -0.7 | -10.2 | -7.4 |
| | N | 9 | 9 | 9 | 9 |
| Total ear weight per plant (g) | $\mu$ | 3.9 | 3.7 | 4.0 | 4.6 |
| | SD | 1.2 | 1.4 | 1.0 | 1.2 |
| | SE | 0.2 | 0.2 | 0.2 | 0.2 |
| | p value | | 0.420 | 0.417 | 0.00762 |
| | Percentage | | -6.2 | 1.4 | 17.8 |
| | N | 36 | 36 | 32 | 36 |
| Seed number per plant | $\mu$ | 55 | 49 | 58 | 61 |
| | SD | 13 | 16 | 13 | 14 |
| | SE | 2 | 3 | 2 | 2 |
| | p value | | 0.0844 | 0.135 | 0.0222 |
| | Percentage | | -10.9 | 6.4 | 11.7 |
| | N | 36 | 36 | 32 | 36 |
| Height (cm) | $\mu$ | 39.6 | 38.6 | 37.6 | 40.5 |
| | SD | 3 | 4 | 5 | 5 |
| | SE | 0.6 | 0.7 | 0.8 | 0.8 |
| | p value | | 0.264 | 0.0267 | 0.175 |
| | Percentage | | -2.6 | -5.0 | 2.4 |
| | N | 36 | 37 | 32 | 36 |
| Average weight per seed (g) | $\mu$ | 0.073 | 0.076 | 0.069 | 0.076 |
| | SD | 0.02 | 0.02 | 0.01 | 0.01 |
| | SE | 0.003 | 0.003 | 0.002 | 0.002 |
| | p value | 0.572 | 0.134 | 0.295 | 0.022 |
| | Percentage | | 3.5 | -6.4 | 2.9 |
| | N | 36 | 36 | 32 | 36 |

***Table S2:*** *Statistical values from the physiology studies. $\mu$ is the mean, SD is the standard deviation, SE is the error on the mean, the p-value is compared to the control, N is the number of repeats, and percentage is the percentage difference from the control*
